# Harmonisation of Physical Activity Variables by Indirect Validation: A Doubly Labelled Water Study

**DOI:** 10.1101/501718

**Authors:** Matthew Pearce, Tom R.P. Bishop, Stephen Sharp, Kate Westgate, Michelle Venables, Nicholas J. Wareham, Søren Brage

## Abstract

Harmonisation of data for pooled analysis relies on the principle of inferential equivalence between variables from different sources. Ideally, this is achieved using models of the direct relationship with gold standard criterion measures, but the necessary validation data are often unavailable. This study examines an alternative method of harmonisation by indirect validation. Starting methods were self-report or accelerometry, from which we derived indirect models of relationships with doubly labelled water (DLW)-measured physical activity energy expenditure (PAEE) using sets of two bridge equations via one of three intermediate measures. Coefficients and performance of indirect models were compared to corresponding direct validation models (linear regression of DLW-measured PAEE on starting methods). Indirect model beta coefficients were attenuated compared to direct model betas (10-63%), narrowing the range of PAEE values; attenuation was greater when bridge equations were weak. Directly and indirectly harmonised models had similar error variance but most indirectly derived values were biased at group-level. Correlations with DLW-measured PAEE were identical after harmonisation using continuous linear but not categorical models. Wrist acceleration harmonised to DLW-measured PAEE via combined accelerometry and heart rate sensing had lowest error variance (24.5%) and non-significant mean bias 0.9 (95%CI: −1.6; 3.4) kJ•day^−1^•kg^−1^. Associations between PAEE and BMI were similar for directly and indirectly harmonised values, but most fell outside the confidence interval of the criterion PAEE-to-BMI association. Indirect models can be used for harmonisation. Performance depends on the measurement properties of original data, variance explained by available bridge equations, and similarity of population characteristics.

---

Harmonisation of exposure and outcome variables is an essential step when integrating different sources of data for the same analysis, such as in meta-analysis of published results, pooled or federated meta-analysis of individual-level data, and global surveillance of risk factors for disease. Analyses of this nature which use information from multiple sources are often constrained by the quality and compatibility of the original data (Fortier et al., 2010). Harmonisation aims to bring together various types and levels of data which represent the same underlying construct (e.g. physical activity, energy intake, body fat percentage etc.) in order to achieve compatibility when methods vary between studies or study phases (Granda & Blasczyk, 2011). The process does not strictly require that precisely the same original collection and processing methods are employed in each study (Fortier et al., 2010), but the harmonised data should be “inferentially equivalent”, i.e. their format, function and meaning are the same (Atkin et al., 2017). This inferential equivalence will depend upon the scientific context and the type of analysis being undertaken.

A common harmonisation approach is conversion to the level of the least detailed information, for example transformation of continuous data to a binary categorisation of low vs. high physical activity level (Kilpelainen et al., 2011). However, this approach loses the resolution of the more detailed data, and may therefore limit the power and scope of subsequent analyses. It is also unclear how well variables harmonised in this way relate to the underlying true latent value of the exposure. An alternative approach to harmonisation is to restrict analyses to only those studies which have assessed and expressed the exposure and outcome in the desired way. This maintains the detail of the contributing data, but – as highlighted by Aune, Norat, Leitzmann, Tonstad, and Vatten (2015) – greatly reduces the proportion of the available data that can be included in evidence synthesis. At best, this leads to loss of power. At worst, this leads to bias if the studies that are included with optimal data have specific characteristics.

Another approach to harmonisation is to use validation studies which report the *direct* statistical (e.g., regression) models of relationships between values from the less precise methods and the latent true level of exposure, as assessed by a construct-specific gold-standard criterion method. A direct validation model permits transformation of original data to the desired harmonised format. The problem is that this mapping approach is often not possible because the ideal validation study employing gold-standard criterion methods has either not been conducted, does not report the statistical model of transformation, or is not applicable to the population or setting in question. This limitation may be more common in particular populations or settings, such as those in which the feasibility or cost of gold- (or even silver-) standard methods is prohibitive. Consequently, some populations and settings may be studied with unsatisfactorily harmonised data or excluded from analyses altogether.

When the ideal direct validation model is unavailable, a potential solution may be to use *indirect* validation models, whereby estimates from the less precise but more feasible field method are mapped to a gold standard criterion using two “bridge equations” which form an indirect route to the criterion via a third intermediate method (Figure 1). This concept is analogous to network meta-analysis in which multiple comparisons can be inferred despite not being directly tested (Lu & Ades, 2004). Indirect harmonisation adopts similar principles in that it utilises existing, ideally published, validity data to derive a new indirect validation model which can be used to transform data without the requirement to conduct de novo fieldwork. By utilising a combination of published bridge equations and existing datasets, this study examines such an approach by comparing the inferential equivalence of data harmonised to the gold standard format via both direct and indirect validation models.

**Figure 1.**
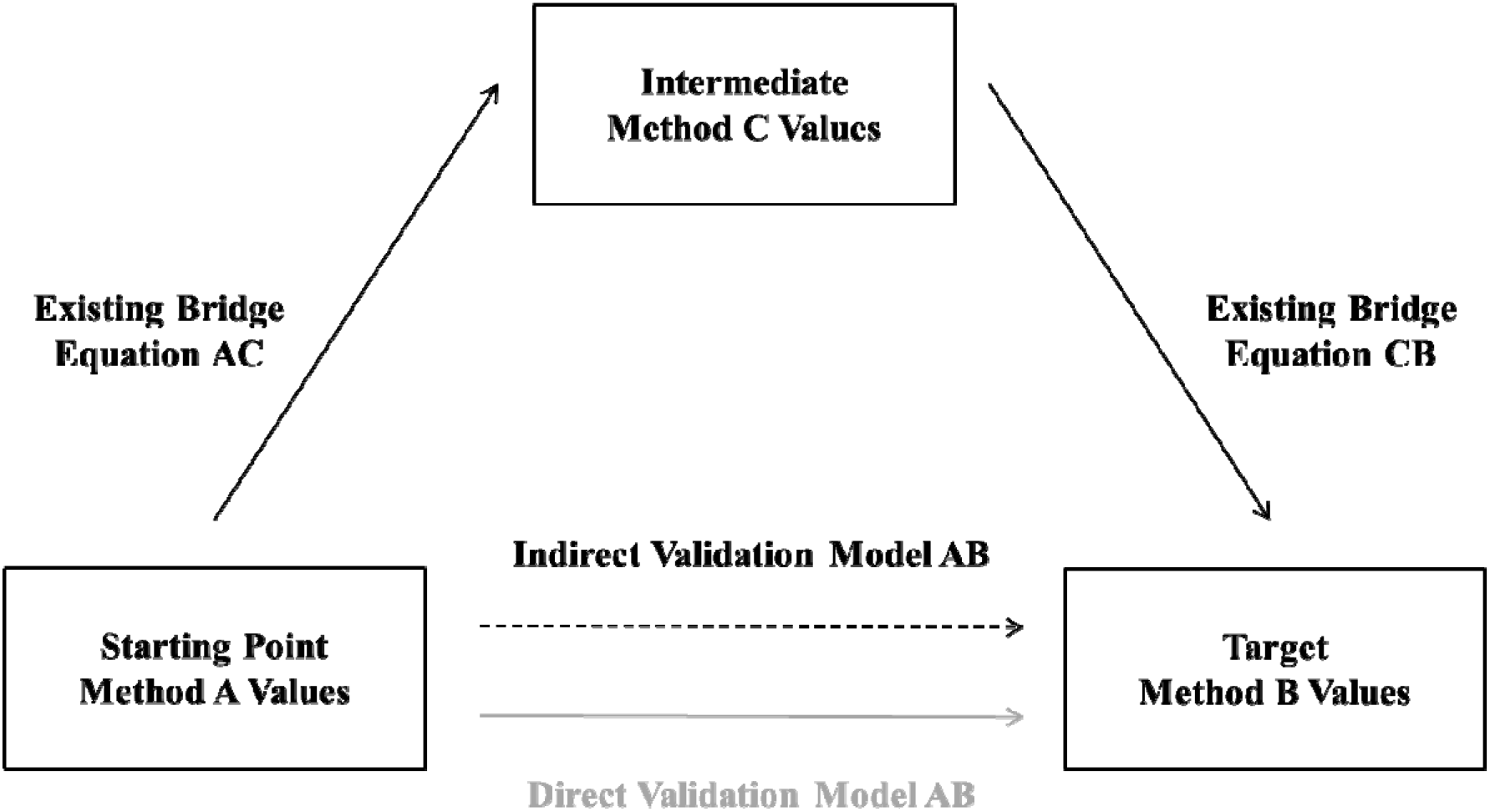
Indirect validation model of the relationship between values from starting point Method A and target Method B via intermediate Method C (broken black arrow). Intermediate methods (Method C) are characterised by already established (published) relationships with both the target criterion method (Method B) and the starting point method (Method A) as indicated by the solid black arrows. The new indirect validation model AB is evaluated against the direct validation model AB (solid grey arrow).

## Methods

### Theoretical modelling framework

We present a hypothetical analysis task requiring harmonisation of, at-first-glance, incompatible starting values from a Method A to the format of values arising from a target criterion Method B. One could use direct validation between Method A and Method B to complete this task (direct validation model AB) but as mentioned above often an alternative approach is needed. Here, we use intermediary values from a third method, Method C, for which separate links to values from Method A and Method B are available. The relationships between values from Method A and Method C (“Bridge Equation AC”) and between values from Method C and Method B (“Bridge Equation CB”) are used to derive the indirect validation model AB as outlined in Figure 1. In order to assess the validity of this indirect validation approach, the indirect model for the AB relationship is compared to the direct model.

### Specific harmonisation task

Here, we use the example of harmonising values from four variations on Method A (the starting point method) to total daily physical activity energy expenditure (PAEE) expressed in kJ•day^−1^•kg^−1^ as measured using the same gold standard criterion (Method B). For simplicity, we will use linear models to describe the links between methods.

The four sets of starting data from Method A were: (1) duration (minutes per day) of moderate to vigorous physical activity (MVPA) derived from the Recent Physical Activity Questionnaire (RPAQ) (Besson, Brage, Jakes, Ekelund, & Wareham, 2010); (2) total daily PAEE expressed in kJ•day^−1^•kg^−1^ derived from RPAQ; (3) the four-level categorical Cambridge Index derived by combining occupational physical activity with time participating in cycling and other physical exercise, giving categories of inactive, moderately inactive, moderately active, and active (Golubic et al., 2014; Peters et al., 2012; Wareham et al., 2003); and (4) the mean daily high-pass filter vector magnitude signal from non-dominant wrist acceleration expressed in milli-g (ACC_WRIST_) (White et al., 2018; White, Westgate, Wareham, & Brage, 2016).

The gold-standard (Method B) for assessing PAEE (kJ•day^−1^•kg^−1^) was the difference between total and resting energy expenditure as measured by the DLW method and two lab-based assessments of resting metabolic rate, coupled with allowance for the diet-induced thermogenic effect.

To derive the indirect validation model between values from the starting point Method A and the criterion Method B, values from one of three variations on the intermediate Method C were used: (1) mean daily trunk acceleration in m•s^−2^ (ACC_TRUNK_) (Brage, Brage, Franks, Ekelund, & Wareham, 2005); (2) total daily PAEE in kJ•day^−1^•kg^−1^ derived from the individually calibrated flex heart rate method (HR) (Brage et al., 2007; Spurr et al., 1988); (3) total daily PAEE in kJ•day^−1^•kg^−1^ from combined ACC_TRUNK_ and HR (ACCHR) (Brage et al., 2004, 2007).

### Data sources for bridge equations

To examine different aspects of the performance of harmonisation using indirect validation models, one of seven variations on Bridge Equation AC was combined with one of three variations on Bridge Equation CB.

If available, we used published equations to derive indirect validation models for harmonisation purposes. If relevant equations were unavailable but correlation coefficients and basic (mean and SD) summary statistics were, we derived the equations through back-transformation of standardised coefficients using the corr2data STATA command. If this was not possible, we used individual-level data from existing datasets to derive equations (pretending these were published validation studies), and subsequently used them alongside existing bridge equations sourced from published work. The following sections describe the data sources of each of the bridge equations.

#### Bridge Equation AC

Five variations on Bridge Equation AC were derived from the Fenland Study, a population-based cohort study of 12,435 adults born between 1950 and 1975 and registered with general practices in Cambridgeshire, United Kingdom. We randomly split this dataset into five subsamples to represent five different and independent validation studies. Participants attended our research facility and completed RPAQ (Besson et al., 2010) and underwent treadmill testing for individual calibration (Brage et al., 2007) whilst fitted with a chest-worn combined heart rate and movement sensor (Actiheart, CamNtech Ltd, Papworth, UK) (Brage et al., 2005). At the end of the clinical assessment, they were instructed to wear this device continuously for six days and nights and carry on with their normal behaviours. Data from RPAQ were used to derive duration (minutes per day) of MVPA by summing duration reported participating in activities with intensity > 3.0 METs (Ainsworth et al., 2011), estimates of PAEE calculated as frequency * duration * intensity (Besson et al., 2010), and the four-level Cambridge Index (Golubic et al., 2014; Wareham et al., 2003). The ACC_TRUNK_ signal from the combined heart and movement sensor was used in the format of mean daily trunk acceleration in m•s^−2^, while the HR signal together with treadmill test data was used to derive an individually calibrated estimate of PAEE as previously described (Brage et al., 2007, 2015). The two signals were also combined (ACCHR) to predict PAEE using branched equation modelling (Brage et al., 2004). The Fenland Study was approved by the Health Research Authority National Research Ethics Service Committee East of England-Cambridge Central, and participants provided written informed consent. The five variations on Bridge Equation AC using self-report as the starting point were derived from the linear regression of: 1) ACC_TRUNK_ on RPAQ MVPA; (2) HR PAEE on RPAQ MVPA; (3) ACCHR PAEE on RPAQ MVPA; (4) ACCHR PAEE on RPAQ PAEE; (5a) ACCHR PAEE on RPAQ Cambridge Index. To examine whether indirect harmonisation is robust to variations in measurement protocol and population (i.e. a non-ideal Bridge Equation), we used an additional linear regression of (5b) ACCHR PAEE on RPAQ Cambridge Index. This was derived from the similar short European Prospective Investigation into Cancer and Nutrition Study Physical Activity Questionnaire (short EPIC-PAQ) administered in the EPIC cohort across 10 European countries (Peters et al., 2012; Wareham et al., 2003) denoted by ACCHR_EUROPE_.

To contrast the harmonisation process of self-report measures as starting points with that of an objective starting measure, two additional AC bridge equations derived from the linear regression of: (6) ACC_TRUNK_ on ACC_WRIST_; and (7) ACCHR PAEE on ACC_WRIST_, which were obtained from published work (White et al., 2016).

#### Bridge Equation CB

Three bridge equations were obtained from published data (Brage et al., 2015) using the linear regression of DLW method total daily PAEE expressed in kJ•day^−1^•kg^−1^ on: (1) ACC_TRUNK_; (2) HR PAEE; and (3) ACCHR PAEE, in doing so linking to the prediction output from a Bridge Equation AC described above.

### Direct Validation Model AB

The inferential equivalence of harmonised PAEE values was assessed using gold standard DLW method PAEE values from the UK Biobank Validation Study (BBVS) reported in detail elsewhere (White et al., 2018). Briefly, rate of carbon dioxide production (rCO_2_) was measured using the ten day DLW method of Schoeller et al. (1986) and converted to total energy expenditure (TEE) using the energy equivalents of CO2 of Elia and Livesey (1988) in 100 participants. Resting metabolic rate (RMR) was measured on two separate days during clinic visits with a fifteen-minute rest test by respired gas analysis (OxyconPro, Jaeger, Germany), and scaled by a factor of 0.94 to account for RMR measurements being conducted in the afternoon rather than the morning (Haugen, Melanson, Tran, Kearney, & Hill, 2003). The closest measurement value (visit 1, visit 2, or their mean) by proximity to the within-person median of predictions of RMR using three equations (Henry, 2005; Nielsen et al., 2000; Watson et al., 2014) was used in analysis. Total daily REE was calculated, with an additional adjustment of sleeping metabolic rate being 5% lower than awake resting metabolic rate (Goldberg, Prentice, Davies, & Murgatroyd, 1988). Diet-induced thermogenesis was estimated using macronutrient intake assessed by food frequency questionnaire as previously described (Brage et al., 2015; Jequier, 2002). The REE and diet-induced thermogenesis were subtracted from TEE and divided by body mass yielding an estimate of total daily PAEE in kJ•day^−1^•kg^−1^.

The four variations on the starting data (3 self-report and 1 objective measure) in the Fenland Study were replicated in BBVS so that four corresponding direct validation models predicting DLW method PAEE could be derived. Participants completed RPAQ and the raw data were used to derive duration of MVPA, PAEE, and the four-level Cambridge Index. In addition, participants were fitted with a tri-axial accelerometer (AX3, Axivity, Newcastle, UK) on the wrist for 9 days and nights whilst continuing with their usual activities. The ACC_WRIST_ signal was used to approximate acceleration as a result of human movement and expressed in milli-g (van Hees et al., 2013). Ethical approval for the study was obtained from Cambridge University Human Biology Research Ethics Committee (Ref: HBREC/2015.16). All participants provided written informed consent.

### Deriving Indirect Validation Model AB

Beta and alpha coefficients for each Indirect Validation Model AB were derived by substituting Bridge Equation AC (eq1) into Bridge Equation CB (eq2) to give formula eq3:

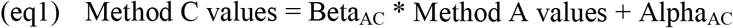

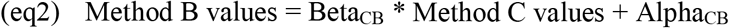

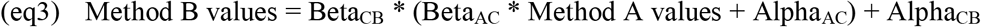

Formula 3 simplifies to give the following formulae for deriving the new alpha (eq3a) and beta (eq3b) coefficients:

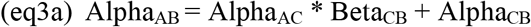

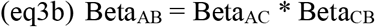

These formulae provide parameter estimates for the Indirect Validation Model AB coefficients but not their standard errors. To propagate the uncertainty in the parameter estimates from Bridge Equation AC and Bridge Equation CB to the new Indirect Validation Model AB, 10 000 values of each parameter were sampled from a normal distribution with mean equal to the observed parameter estimate and standard deviation equal to the standard error of that parameter estimate; the formulae above were then applied to the sampled values. The means and standard deviations of the resulting distributions for BetaAB and AlphaAB were used as the coefficient point estimates and standard errors, respectively.

For the indirect validation model using the categorical Cambridge Index derived from RPAQ, the categorical data were replaced by one constant and three dummy variables to represent four levels. The above steps were then applied in the same way, but repeated for each of the four values of Beta_AC_.

We meta-analysed the newly derived beta and alpha coefficients from each of three indirect validation models predicting PAEE from duration of MVPA, thus generating a fifth indirect combined prediction equation using all available information; this represents the scenario where harmonisation is performed using multiple validation studies of the same instrument.

### Analysis

The inferential equivalence of each permutation of Indirect Validation Model AB was assessed alongside an equivalent Direct Validation Model AB derived from the linear regression of individual-level Method B data on Method A data available for 100 participants in the BBVS. Method B values (PAEE) were predicted from Method A values using each of the direct and indirect validation models and compared with values from the observed criterion PAEE (i.e. the “true” PAEE exposure using the DLW method) by calculating the mean bias and 95% limits of agreement, root mean square error (RMSE), and Spearman correlation to assess the similarity with which individuals were ranked. Note in this evaluation scenario, the mean bias of directly mapped relationships is always zero. We derived the theoretical combined explained variance as the product of the r^2^ values from the two linear bridge equations for each Indirect Validation Model.

Finally, to demonstrate utility, we examined the associations between all PAEE estimates and body mass index (BMI) using multivariable linear regression adjusted for age and sex in a subset of 1695 participants in the Fenland Study. All data processing and analyses were performed in STATA/SE 14.2 (StataCorp, TX, USA).

## Results

The characteristics of participants from each of the sources of data are described in Table 1. The participants in the Brage et al. (2015) study were younger and more active with lower BMI than participants in BBVS and the Fenland Study, including the subset reported in White et al. (2016). Participants in EPIC were less active than those in the Fenland Study. The criterion value of PAEE from the DLW method in the BBVS had a mean (SD) of 49.7 (16.2) kJ •kg^−1^•day^−1^ and a range of 8.6 to 90.8 kJ•kg^−1^•day^−1^.

**Table 1.** Participant haracteristics.

|  | Fenland Study | White et al.<br>(2016) | Peters et al.<br>(2012) | Brage et al.<br>(2015) | BBVS |
| --- | --- | --- | --- | --- | --- |
| N | 10 602 | 1050 | 1941 | 46 | 100 |
| Age (years) | 48 (7) | 50 (7) | 53 (8) | 34 (9) | 54 (7) |
| Female (%) | 52 | 48 | 70 | 50 | 50 |
| BMI ($\text{kg}\cdot\text{m}^{-2}$) | 27 (5) | 26 (4) | 26 (4) | 25 (3) | 27 (3) |
| $\text{ACC}_{\text{TRUNK}}$ ($\text{m}\cdot\text{s}^{-2}$) | .126 (.055) | .125 (.055) | - | .237 (.090) | - |
| HR PAEE ($\text{kJ}\cdot\text{day}^{-1}\cdot\text{kg}^{-1}$) | 71 (43) | - | - | 67 (42) | - |
| ACCHR PAEE ($\text{kJ}\cdot\text{day}^{-1}\cdot\text{kg}^{-1}$) | 55 (22) | - | 44 (16) | 69 (25) | - |
| RPAQ MVPA (minutes$\cdot\text{day}^{-1}$) | 112 (152) | - | - | - | 128 (146) |
| RPAQ PAEE ($\text{kJ}\cdot\text{day}^{-1}\cdot\text{kg}^{-1}$) | 38 (32) | - | - | - | 40 (29) |
| CAM Inactive (%) | 13 | - | 15 | - | 7 |
| CAM Moderately inactive (%) | 54 | - | 33 | - | 37 |
| CAM Moderately active (%) | 28 | - | 28 | - | 30 |
| CAM Active (%) | 17 | - | 24 | - | 26 |
| DLW method PAEE ($\text{kJ}\cdot\text{day}^{-1}\cdot\text{kg}^{-1}$) | - | - | - | 66 (24) | 50 (16) |
| $\text{ACC}_{\text{WRIST}}$ (milli-g) | - | 48 (11) | - | - | 43 (10)* |

Abbreviations: $\text{ACC}_{\text{TRUNK}}$ = mean daily trunk acceleration; $\text{ACC}_{\text{WRIST}}$ = mean daily high pass filter vector magnitude wrist acceleration signal; ACCHR = combined heart rate and movement sensing; BBVS = Biobank Validation Study; BMI = body mass index; CAM = Cambridge Index; DLW = doubly labelled water; HR = heart rate; MVPA = moderate-to-vigorous physical activity; PAEE = physical activity energy expenditure; RPAQ = Recent Physical Activity Questionnaire.
Values are mean (SD) unless specified otherwise.
\*n = 97.

The combinations of Bridge Equation AC and Bridge Equation CB used to generate the Indirect Validation Model AB are shown in Table 2 alongside their r^2^ values. The newly derived indirect validation models are plotted alongside the direct validation models in Figure 2.

**Table 2.** Bridge equations used to derive ndirect validation models.

| Bridge equation | Starting Point Value A | Intermediate Value C | Target Value B | N | $\beta$ (SE) | $\alpha$ (SE) | $r^2$ |
| --- | --- | --- | --- | --- | --- | --- | --- |
| **Indirect harmonisation of RPAQ MVPA via ACC<sub>TRUNK</sub>** |  |  |  |  |  |  |  |
| AC | RPAQ MVPA (minutes•day<sup>-1</sup>) | ACC<sub>TRUNK</sub> (m•s<sup>-2</sup>) | - | 2121 | 5.84•10<sup>-5</sup> (7.9•10<sup>-6</sup>) | .1199 (.0015) | .02 |
| CB | - | ACC<sub>TRUNK</sub> (m•s<sup>-2</sup>) | DLW PAEE (kJ•day<sup>-1</sup>•kg<sup>-1</sup>) | 46 | 165 (32) | 26.7 (8.2) | .37 |
| **Indirect harmonisation of RPAQ MVPA via PAEE from HR** |  |  |  |  |  |  |  |
| AC | RPAQ MVPA (minutes•day<sup>-1</sup>) | HR PAEE (kJ•day<sup>-1</sup>•kg<sup>-1</sup>) | - | 2121 | .0840 (.0061) | 60.9 (1.2) | .08 |
| CB | - | HR PAEE (kJ•day<sup>-1</sup>•kg<sup>-1</sup>) | DLW PAEE (kJ•day<sup>-1</sup>•kg<sup>-1</sup>) | 46 | .34 (.07) | 42.7 (5.8) | .34 |
| **Indirect harmonisation of RPAQ MVPA via PAEE from ACCHR** |  |  |  |  |  |  |  |
| AC | RPAQ MVPA (minutes•day<sup>-1</sup>) | ACCHR PAEE (kJ•day<sup>-1</sup>•kg<sup>-1</sup>) | - | 2120 | .0390 (.0030) | 50.69 (.57) | .07 |
| CB | - | ACCHR PAEE (kJ•day<sup>-1</sup>•kg<sup>-1</sup>) | DLW PAEE (kJ•day<sup>-1</sup>•kg<sup>-1</sup>) | 46 | .66 (.11) | 20.0 (8.1) | .45 |
| **Indirect harmonisation of RPAQ PAEE via PAEE from ACCHR** |  |  |  |  |  |  |  |
| AC | RPAQ PAEE (kJ•day<sup>-1</sup>•kg<sup>-1</sup>) | ACCHR PAEE (kJ•day<sup>-1</sup>•kg<sup>-1</sup>) | - | 2120 | .239 (.014) | 45.63 (.69) | .12 |
| CB | - | ACCHR PAEE (kJ•day<sup>-1</sup>•kg<sup>-1</sup>) | DLW PAEE (kJ•day<sup>-1</sup>•kg<sup>-1</sup>) | 46 | .66 (.11) | 20.0 (8.1) | .45 |
| **Indirect harmonisation of Cambridge Index via PAEE from ACCHR** |  |  |  |  |  |  |  |
| AC | RPAQ Cambridge Index | ACCHR PAEE (kJ•day<sup>-1</sup>•kg<sup>-1</sup>) | - | 2120 | *Inactive =0;<br>Moderately inactive = 5.6 (1.5);<br>Moderately active = 13.0 (1.6);<br>Active = 24.2 (1.7) | 44.2 (1.3) | .12 |
| CB | - | ACCHR PAEE (kJ•day<sup>-1</sup>•kg<sup>-1</sup>) | DLW PAEE (kJ•day<sup>-1</sup>•kg<sup>-1</sup>) | 46 | .66 (.11) | 20.0 (8.1) | .45 |
| **Indirect harmonisation of Cambridge Index via PAEE from ACCHR<sub>EUROPE</sub>** |  |  |  |  |  |  |  |
| AC | RPAQ Cambridge Index | ACCHR<sub>EUROPE</sub> PAEE (kJ•day<sup>-1</sup>•kg<sup>-1</sup>) | - | 1941 | *Inactive =0;<br>Moderately inactive = 4.6 (1.1);<br>Moderately active = 9.1 (1.1);<br>Active = 14.8 (1.1) | 36.14 (.88) | .10 |
| CB | - | ACCHR<sub>EUROPE</sub> PAEE (kJ•day<sup>-1</sup>•kg<sup>-1</sup>) | DLW PAEE (kJ•day<sup>-1</sup>•kg<sup>-1</sup>) | 46 | .66 (.11) | 20.0 (8.1) | .45 |
| **Indirect harmonisation of ACC<sub>WRIST</sub> via ACC<sub>TRUNK</sub>** |  |  |  |  |  |  |  |
| AC | ACC<sub>WRIST</sub> (milli-g) | ACC<sub>TRUNK</sub> (m•s<sup>-2</sup>) | - | 1050 | 4.78•10<sup>-3</sup> (9.0•10<sup>-5</sup>) | -.097 (.0036) | .53 |
| CB | - | ACC<sub>TRUNK</sub> (m•s<sup>-2</sup>) | DLW PAEE (kJ•day<sup>-1</sup>•kg<sup>-1</sup>) | 46 | 165 (32) | 26.7 (8.2) | .37 |
| **Indirect harmonisation of ACC<sub>WRIST</sub> via PAEE from ACCHR** |  |  |  |  |  |  |  |
| AC | ACC<sub>WRIST</sub> (milli-g) | ACCHR PAEE (kJ•day<sup>-1</sup>•kg<sup>-1</sup>) | - | 1050 | 1.232 (.012) | -6.90 (.45) | .67 |
| CB | - | ACCHR PAEE (kJ•day<sup>-1</sup>•kg<sup>-1</sup>) | DLW PAEE (kJ•day<sup>-1</sup>•kg<sup>-1</sup>) | 46 | .66 (.11) | 20.0 (8.1) | .45 |

\*Constant and three dummy variables used to represent four-level Cambridge Index.
Abbreviations: ACC<sub>TRUNK</sub> = mean daily trunk acceleration; ACC<sub>WRIST</sub> = mean daily high pass filter vector magnitude wrist acceleration signal; ACCHR = combined heart rate and movement sensing; ACCHR<sub>EUROPE</sub> = combined heart rate and movement sensing from European population; DLW = doubly labelled water method; HR = heart rate; MVPA = moderate-to-vigorous physical activity; PAEE = physical activity energy expenditure; RPAQ = Recent Physical Activity Questionnaire; SE = standard error.

**Figure 2.**
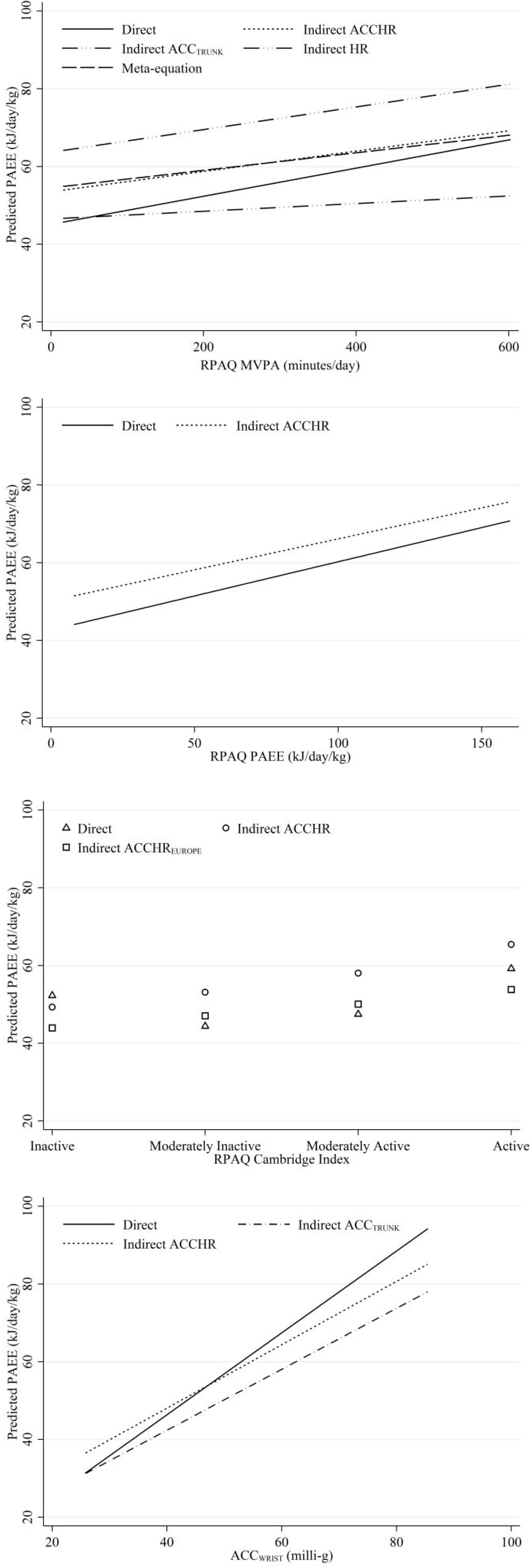
Comparison of direct and indirect harmonisation models by starting data type. *Abbreviations:* ACC_TRUNK_ = mean daily trunk acceleration; ACCHR = combined heart rate and movement sensing; ACCHR_EUROPE_ = combined heart rate and movement sensing from European population; ACC_WRIST_ = mean daily high pass filter vector magnitude wrist acceleration signal; HR = heart rate; MVPA = moderate-to-vigorous physical activity; PAEE = physical activity energy expenditure; RPAQ = Recent Physical Activity Questionnaire.

Table 3 reports the coefficients and performance of validation models predict DLW method PAEE (kJ•kg^−1^•day^−1^) from a continuous duration estimate of MVPA (minutes•day^−1^) derived from RPAQ. The beta coefficients for indirect validation models were attenuated compared with the direct validation model beta (also see Figure 2). The correlation between the predicted and DLW method PAEE values was preserved irrespective of harmonisation method; however at group-level the values from ACCHR and HR indirect validation models were significantly biased. Each validation model was characterised by a narrowing of the range of PAEE values, and this was particularly pronounced when using ACC_TRUNK_ values as the intermediate method, the indirect validation model with the smallest combined explained variance, most attenuated beta coefficient, and widest limits of agreement. The HR indirect validation model resulted in predictions with the largest RMSE while the ACC_TRUNK_ and ACCHR indirect validation model predictions were similarly precise. Coefficients generated by meta-analysing the three indirect validation models had smaller standard errors and performance was similar to that of harmonisation via ACCHR.

**Table 3.** Model coefficients and performance for predicting DLW method PAEE from RPAQ MVPA.

|  | Direct<br>BBVS | Indirect<br>via ACC<sub>TRUNK</sub> | Indirect<br>via HR | Indirect<br>via ACCHR | Meta-equation |
| --- | --- | --- | --- | --- | --- |
| **Coefficients** |  |  |  |  |  |
| $\beta$ (SE) | .036 (.011) | .0099 (.0032) | .0291 (.0081) | .0261 (.0062) | .0225 (.0012) |
| $\alpha$ (SE) | 45.1 (2.1) | 46.5 (12.2) | 63.7 (10.6) | 53.5 (13.9) | 54.5 (.23) |
| Combined explained variance | - | .007 | .027 | .032 | - |
| **Performance vs DLW PAEE** |  |  |  |  |  |
| Mean bias (kJ•day<sup>-1</sup>•kg<sup>-1</sup>)[95% CI] | - | -1.9 [-5.1; 1.2] | 17.7 [14.6; 20.7] | 7.1 [4.1; 10.1] | 7.7 [4.6; 10.7] |
| Mean bias (%) | - | -3.9 | 35.6 | 14.3 | 15.4 |
| Limits of agreement* (kJ•day<sup>-1</sup>•kg<sup>-1</sup>) | - | -34; 30 | -13; 48 | -24; 37 | 23; 39 |
| RMSE (kJ•day<sup>-1</sup>•kg<sup>-1</sup>) | 15.3 | 15.8 | 23.4 | 16.9 | 17.2 |
| RMSE (%) | 30.7 | 32.0 | 47.0 | 34.0 | 34.6 |
| Spearman correlation* (rho) | .37 | .37 | .37 | .37 | .37 |
| Range (kJ•day<sup>-1</sup>•kg<sup>-1</sup>) | 46; 67 | 47; 52 | 64; 81 | 54; 69 | 55; 68 |

Note: Combined explained variance is the product of the explained variance of the two bridge equations. It is not the explained variance ( $r^2$ ) of the indirect model.
Abbreviations: ACC<sub>TRUNK</sub> = mean daily trunk acceleration; ACCHR = combined heart rate and movement sensing; BBVS = Biobank Validation Study; DLW = doubly labelled water; HR = heart rate; MVPA = moderate-to-vigorous physical activity; PAEE = physical activity energy expenditure; RMSE = root mean square error; RPAQ = Recent Physical Activity Questionnaire; SE = standard error.

Table 4 reports the coefficients and performance of validation models predicting DLW method PAEE (kJ•kg^−1^•day^−1^) from a continuous estimate of total daily PAEE (kJ•kg^−1•^day^−1^) derived from RPAQ. Compared with the raw values of RPAQ-derived PAEE, the indirect and direct validation models reduced group-level mean bias and approximately halved RMSE. As for MVPA estimates, the correlation between RPAQ PAEE values and criterion PAEE was maintained regardless of whether values were raw, directly or indirectly harmonised. The ranges of the indirectly and directly harmonised values were much reduced compared with those from the raw RPAQ and those of the criterion PAEE.

**Table 4.** Model coefficients and performance for predicting DLW method PAEE from RPAQ PAEE.

|  | Direct<br>BBVS | Indirect<br>via ACCHR | Non-harmonised<br>RPAQ PAEE |
| --- | --- | --- | --- |
| **Coefficients** |  |  |  |
| $\beta$ (SE) | .176 (.054) | .159 (.035) | - |
| $\alpha$ (SE) | 42.7 (2.7) | 50.2 (13.5) | - |
| Combined explained variance | - | .054 | - |
| **Performance vs DLW PAEE** |  |  |  |
| Mean bias ($\text{kJ}\cdot\text{day}^{-1}\cdot\text{kg}^{-1}$) [95% CI] | - | 6.9 [3.8; 9.9] | -9.6 [-15.2; -4.0] |
| Mean bias (%) | - | 13.7 | -19.3 |
| Limits of agreement ($\text{kJ}\cdot\text{day}^{-1}\cdot\text{kg}^{-1}$) | - | -24; 38 | -66; 47 |
| RMSE ($\text{kJ}\cdot\text{day}^{-1}\cdot\text{kg}^{-1}$) | 15.3 | 16.8 | 29.8 |
| RMSE (%) | 30.8 | 33.8 | 60.0 |
| Spearman correlation ($\rho$) | .38 | .38 | .38 |
| Range ($\text{kJ}\cdot\text{day}^{-1}\cdot\text{kg}^{-1}$) | 44; 71 | 51; 76 | 8; 160 |

*Note:* Combined explained variance is the product of the explained variance of the two bridge equations. It is not the explained variance ( $r^2$ ) of the indirect model.
*Abbreviations:* ACCHR = combined heart rate and movement sensing; BBVS = Biobank Validation Study; DLW = doubly labelled water; PAEE = physical activity energy expenditure; RMSE = root mean square error; RPAQ = Recent Physical Activity Questionnaire; SE = standard error.

Table 5 reports the coefficients and performance of validation models predicting DLW method PAEE (kJ•kg^−1^•day^−1^) using the categorical Cambridge Index derived from RPAQ. Correlations with criterion DLW method PAEE were weaker, and differed for direct and indirect harmonisation. For the direct validation model, the value of PAEE assigned to being “inactive” was greater than that assigned to being “moderately inactive” and “moderately active”; the indirect validation model values of PAEE for each category were ordered more intuitively (also see Figure 2). RMSEs for the indirectly and directly harmonised values were similar, and also similar to RMSEs using continuous RPAQ data described above, however the group-level values were again biased when using the Fenland Study data for Bridge Equation AC. The indirect validation model derived using the nonideal Bridge Equation AC from a less active European population showed unbiased group-level estimates of PAEE.

**Table 5.**
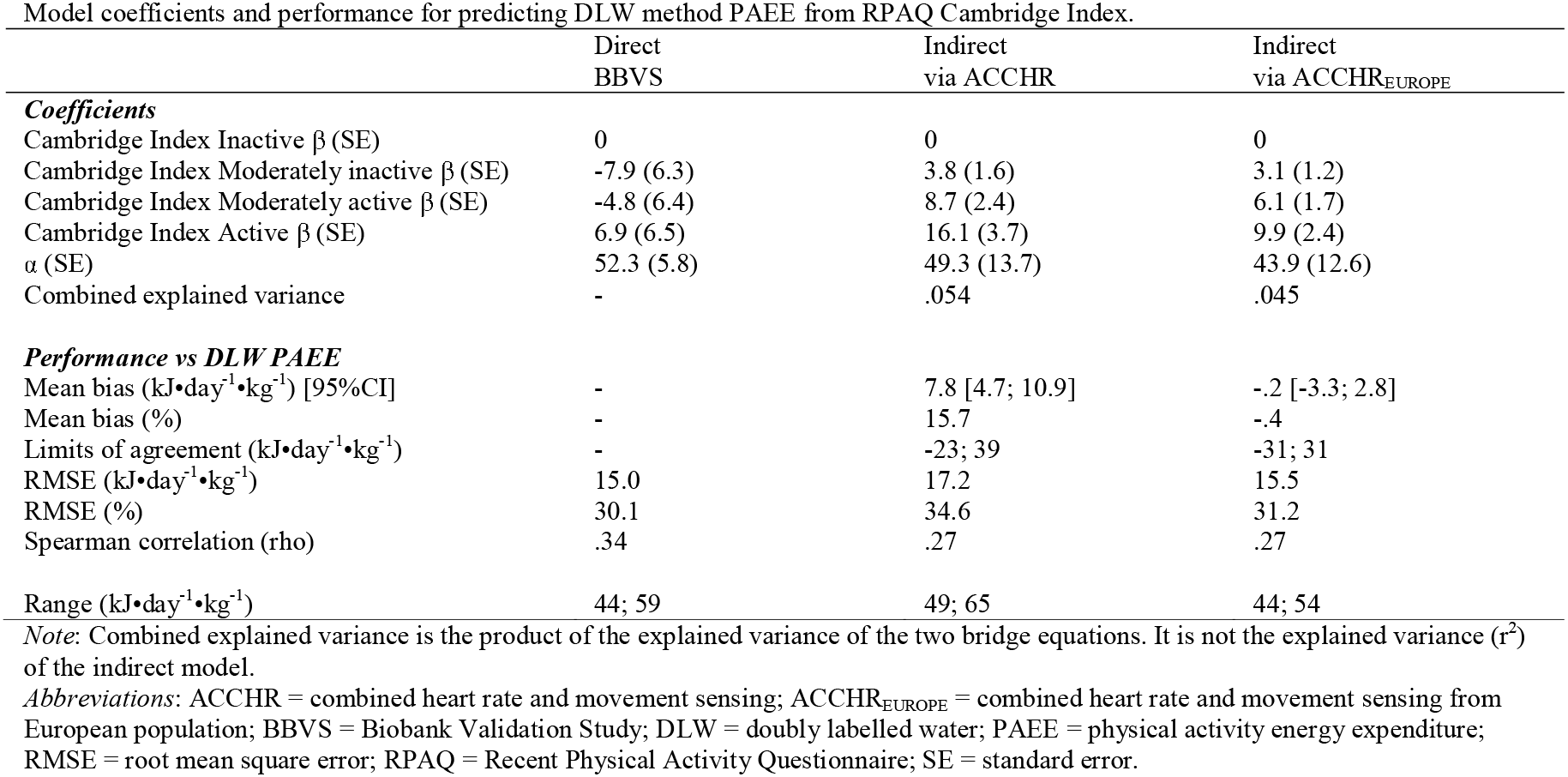
Model coefficients and performance for predicting DLW method PAEE from RPAQ Cambridge Index.

Table 6 reports the coefficients and performance of validation models predicting DLW method PAEE (kJ•kg^−1^•day^−1^) from ACC_WRIST_ (milli-g). Compared with the RPAQ-derived models described above, models based on accelerometer data as starting point and mapping via other objective methods resulted in stronger correlations of estimated PAEE with DLW method PAEE. The RMSEs observed for both the indirect and direct validation models were smaller than for any of the RPAQ validation models, and with higher combined explained variance; the range of PAEE values was also better preserved as shown in Figure 2.

**Table 6.** Model coefficients and performance for predicting DLW method PAEE from ACC_WRIST_.

| | Direct<br>BBVS | Indirect<br>via $ACC_{TRUNK}$ | Indirect<br>via ACCHR |
| --- | --- | --- | --- |
| **Coefficients** |  |  |  |
| $\beta$ (SE) | 1.05 (.12) | .79 (.17) | .81 (.14) |
| $\alpha$ (SE) | 4.2 (5.3) | 10.9 (5.6) | 15.5 (7.6) |
| Combined explained variance | - | .196 | .302 |
| **Performance vs DLW PAEE** |  |  |  |
| Mean bias ($\text{kJ} \cdot \text{day}^{-1} \cdot \text{kg}^{-1}$) [95% CI] | - | -4.8 [-7.2; -2.3] | .9 [-1.6; 3.4] |
| Mean bias (%) | - | -9.5 | 1.8 |
| Limits of agreement ($\text{kJ} \cdot \text{day}^{-1} \cdot \text{kg}^{-1}$) | -24; 24 | -29; 20 | -24; 25 |
| RMSE ($\text{kJ} \cdot \text{day}^{-1} \cdot \text{kg}^{-1}$) | 11.9 | 13.1 | 12.2 |
| RMSE (%) | 24.0 | 26.4 | 24.5 |
| Spearman correlation ($\rho$) | .69 | .69 | .69 |
| Range ($\text{kJ} \cdot \text{day}^{-1} \cdot \text{kg}^{-1}$) | 31; 94 | 31; 78 | 37; 85 |

*Note:* Combined explained variance is the product of the explained variance of the two bridge equations. It is not the explained variance ( $r^2$ ) of the indirect model.
*Abbreviations:* $ACC_{TRUNK}$ = mean daily trunk acceleration; $ACC_{WRIST}$ = mean daily high pass filter vector magnitude wrist acceleration signal; ACCHR = combined heart rate and movement sensing; BBVS = Biobank Validation Study; DLW = doubly labelled water; PAEE = physical activity energy expenditure; RMSE = root mean square error; SE = standard error.

To demonstrate utility of the different harmonisation approaches, we examined the associations of PAEE with BMI in the Fenland study (n=1695); Figure 3 shows the beta (95% confidence intervals) of these linear associations by harmonisation method and starting data format, alongside that from the silver-standard ACCHR PAEE. PAEE estimates harmonised from ACC_WRIST_ showed statistically significant inverse associations with BMI. There was a striking difference between models for the Cambridge Index, with indirectly harmonised PAEE showing statistically significant inverse associations with BMI and the directly harmonised PAEE showing no relationship. Other associations of indirectly harmonised PAEE values with BMI were similar but more uncertain when compared to the direct equivalent. All associations using PAEE harmonised from continuous RPAQ data were weak with 95% confidence intervals crossing zero. MVPA data indirectly harmonised to PAEE via ACCTRUNK values resulted in the widest confidence intervals.

**Figure 3.**
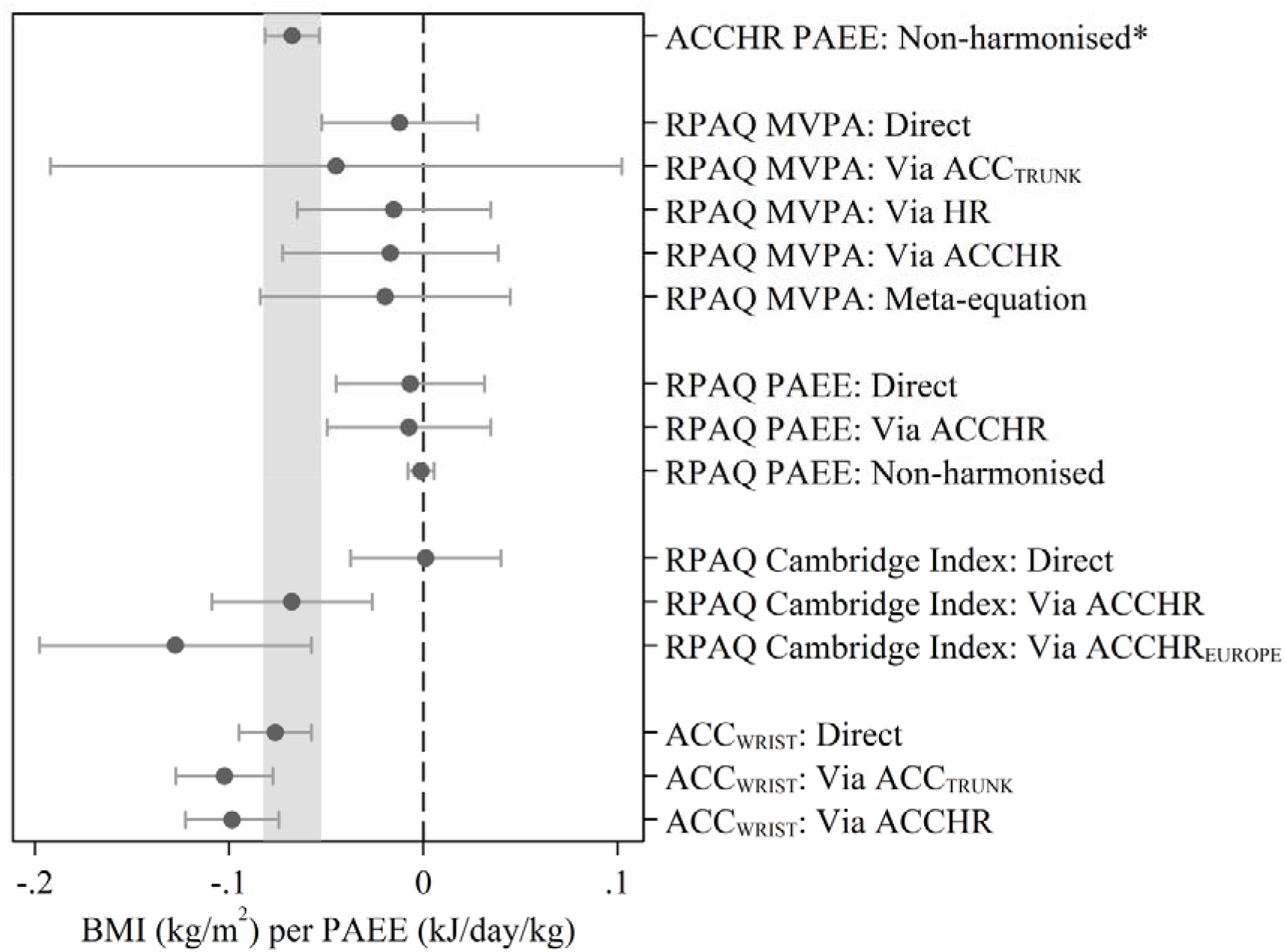
Association (beta coefficients and 95% confidence intervals) between PAEE and BMI, by exposure estimation method (Fenland Study, n=1695 subsample with wrist acceleration). *Note:* Associations are adjusted for age and sex. *Abbreviations*: ACCTRuNK = mean daily trunk acceleration; ACCHR = combined ACC_TRUNK_ and heart rate sensing; ACCHR_EUROPE_ = combined heart rate and movement sensing from European population; BMI = body mass index; ACC_WRIST_ = mean daily high-pass filtered vector magnitude wrist acceleration; HR = heart rate; PAEE = physical activity energy expenditure. *ln the absence of doubly labelled water-assessed PAEE in a large cohort, a silver standard (ACCHR) has been used for comparison for the cross-sectional association.

## Conclusions

This study is the first to examine and evaluate an indirect validation technique for harmonisation, an essential step in any study which combines data from different sources in the same analysis. Our findings indicate that indirect validation models can be employed to harmonise data to a compatible format in the absence of the ideal validation study, but that the harmonised values may still be biased at group-level, have a narrower value range compared to the criterion, and that gains in precision are dependent upon the variance explained by the contributing bridge equations. These findings reinforce the necessity to carefully consider the inferential equivalence of harmonised data and “the truth” according to the scientific context and the purpose of the analyses being performed (Fortier et al., 2010).

For analyses such as exposure-disease associations which primarily rely on relative validity, harmonisation using indirect validation models is beneficial in that it increases precision while retaining the correlation between the original data and estimates of the latent truth from the gold standard criterion (when all bridge equations are linear). Our results for RPAQ-derived PAEE demonstrate that even when data pre-exist in a seemingly compatible format, improvements in precision are possible following harmonisation. Furthermore, even if no harmonisation need be conducted, the newly derived indirect validation model could be used to perform measurement error correction by regression calibration (Keogh & White, 2014).

Using indirect validation models revealed some unexpected advantages. When harmonising Cambridge Index categorical data to a continuous estimate of PAEE, the direct validation model assigned a higher PAEE value to level 1 than both level 2 and 3, whereas the indirectly validation model was more logically ordered by mean PAEE in kJ•kg^−1^•day^−1^ using the indirect relationships in the Fenland sample or the published EPIC data. The illogical ordering likely contributed to the null PAEE-BMI association for directly harmonised PAEE, whereas the two logically ordered indirectly harmonised PAEE estimates both showed inverse relationships. Given that validation studies using comparisons with a gold-standard criterion are often relatively small, mapping of categorical data from questionnaires to continuous metrics may be unduly influenced by incorrectly classified individuals with extreme values. In a larger study such as a prospective cohort like Fenland, the influence of incorrectly classified individuals on category means is diminished resulting in more logically ordered group means. Thus in certain scenarios and under some feasibility constraints, it is possible that indirect validation models using bridge equations sourced from larger studies with silver-standard assessment are actually preferable to harmonisation using a direct validation model from a smaller study using the gold standard. Although indirect harmonisation is a potential solution in the absence of direct validation models, these findings also suggest investigators should make use of all available information – both direct and indirect – to optimise harmonisation efforts.

We observed that beta coefficients of indirect validation models were attenuated compared to those from direct models, and that the level of attenuation is related to the variance explained by the bridge equations being combined. We calculated the theoretical combined explained variance which summarises the r^2^ values of the two contributing bridge equations derived in separate studies; indirect validation models with a higher value tended to have less attenuated beta coefficients, greater precision and less narrowing of the range of resulting values. Shrinking of the range of harmonised exposure values towards the mean when compared to the original data and the criterion will have implications for dose-response analyses which aim to assess the shape of any relationship across the full exposure range. It was noticeable that the indirect validation model for harmonising MVPA to PAEE via ACC_TRUNK_ had the most attenuated beta coefficient resulting in a very narrow range of PAEE values and the widest confidence intervals for the applied example of studying the PAEE association with BMI. A key finding was that associations between PAEE and BMI were similar for data harmonised using both direct and indirect models; the two techniques may therefore be inferentially equivalent but depending upon the validity of the original method, neither may result in data inferentially equivalent to the true level of exposure. The utility of all harmonisation equations is influenced by the quality of the data being harmonised and the error of the methods used in the bridge equations.

For analyses which depend upon absolute agreement between harmonised data and the latent truth, even greater caution is required as most values from indirect harmonisation were biased at group-level. The bias was typically <15% and did not appear to be related to the combined explained variance. The biases observed may therefore reflect differences in the populations from which the bridge equations and the data undergoing harmonisation are sourced. Perhaps most strikingly, the criterion measures of PAEE in the BBVS and Brage et al. (2015) differed on average by 22%. The Brage et al. (2015) participants were younger and had lower BMI so these are likely to be true differences, even though some of the bias observed may be due to minor differences in the criterion method used to estimate PAEE in the two studies, e.g. different RMR protocols. We examined the effect of using alternative Bridge Equations from potentially incompatible measurement tools and populations by deriving two indirect models for harmonising the RPAQ Cambridge Index to PAEE. The indirect model using a Bridge Equation AC derived from the Fenland Study resulted in biased group-level estimates, whereas the ‘non-ideal’ Bridge Equation AC derived from published data using the EPIC-PAQ in the less active European population did not. It is possible that the lower PAEE in this population counteracted any bias resulting from the higher PAEE in the second Bridge Equation from Brage et al. (2015). The contrast may also be partially attributable to differences between RPAQ and EPIC-PAQ. Any incompatibility of the populations and assessment methods from which the two Bridge Equations are derived must therefore be considered alongside the data being harmonised.

Further work is necessary to examine the suitability of combining bridge equations from different populations and the generalisability of the resulting indirect validation models, but our findings suggest that harmonisation is sensitive to differences in the true level of exposure between participants in the bridge equation studies and those in the dataset being harmonised.

One potential solution to population differences could be the addition of covariates to indirect validation models, or, if the sample is large enough, stratification of results by covariates. However, this would require some existing published bridge equations to be re-derived, and limit future harmonisation to datasets with compatible covariates. Moreover, adding covariates to the prediction likely requires that these are always adjusted for in subsequent association analyses, as these may otherwise be driven by the covariates, i.e. confounded. In addition and on a more practical level, it is often the case that regression equations are not published, and this limits harmonisation efforts despite some of the necessary fieldwork having been conducted. We have added beta and alpha coefficients alongside their standard errors of the bridge equations used in the present study to the online repository called the Diet, Anthropometry, and Physical Activity Measurement Toolkit (www.measurement-toolkit.org) which enables sharing of such results and which should increase the discoverability and use of population-specific validation models.

A potential limitation of the current work is that only simple linear bridge equations were combined and that the participants in the studies used were all from the same region of the UK; the generalisability of the technique to other populations therefore remains unclear. While it was a strength of this work that we conducted indirect harmonisation with several different combinations of methods, there are many more methods in use and the viability of this process with other combinations, particularly when both bridge equations have weak r^2^ is difficult to judge. In this study we were able to assess the validity of indirect harmonisation using the equivalent direct relationship from a study with gold-standard measures. This will not be possible in most other scenarios as direct models would be unavailable – the exact problem that indirect harmonisation tries to resolve. In these circumstances it is not possible to assess the precision or mean bias of indirectly harmonised estimates; the variance explained by the two bridge equations provides an indication, but additional research is required to examine this formally.

In summary, indirect validation models can harmonise data to compatible format when direct validation models are not available, and can therefore improve the inclusivity and resolution of data in analyses integrating information from different sources. The process preserves correlations for linearly mapped continuous variables and increases precision, however the range of exposure values is narrowed, and each of these traits is reliant on the quality of the original data and the variance explained by available bridge equations. Further work is required to examine the sources of bias and address potential difficulties when generalising population specific equations.

## Acknowledgements

We are indebted to the volunteers who took part in the Fenland Study and the Biobank Validation Study. We thank the MRC Epidemiology Unit functional group teams for study co-ordination, data collection, IT and data management in this study, as well as the principal investigators of the Fenland Study and the Biobank Validation Study. In particular we would like to thank Tom White, Stefanie Hollidge and Lewis Griffiths for assistance with physical activity data processing, and Eirini Trichia from the MRC Epidemiology Unit for processing the FFQ data with the FETA package. We would also like to thank the stable isotope team from the MRC Elsie Widdowson Laboratory: Priya Singh, Elise Orford and Kevin Donkers for the DLW preparation and analysis. David Vaughan from the MRC Epidemiology Unit is acknowledged for his assistance in creating the instrument library on www.measurement-toolkit.org.

